# Ensemble-GNN: federated ensemble learning with graph neural networks for disease module discovery and classification

**DOI:** 10.1101/2023.03.22.533772

**Authors:** Bastian Pfeifer, Hryhorii Chereda, Roman Martin, Anna Saranti, Sandra Clemens, Anne-Christin Hauschild, Tim Beißbarth, Andreas Holzinger, Dominik Heider

**Affiliations:** Institute for Medical Informatics, Statistics and Documentation. Medical University Graz, Austria; Medical Bioinformatics, University Medical Center Göttingen, Göttingen, Germany; Data Science in Biomedicine, Department of Mathematics and Computer Science, University of Marburg, Marburg, Germany; Human-Centered AI Lab, University of Natural Resources and Life Sciences, Vienna, Austria; Institute for Medical Informatics, University Medical Center Göttingen, Göttingen, Germany

## Abstract

Federated learning enables collaboration in medicine, where data is scattered across multiple centers without the need to aggregate the data in a central cloud. While, in general, machine learning models can be applied to a wide range of data types, graph neural networks (GNNs) are particularly developed for graphs, which are very common in the biomedical domain. For instance, a patient can be represented by a protein-protein interaction (PPI) network where the nodes contain the patient-specific omics features. Here, we present our Ensemble-GNN software package, which can be used to deploy federated, ensemble-based GNNs in Python. Ensemble-GNN allows to quickly build predictive models utilizing PPI networks consisting of various node features such as gene expression and/or DNA methylation. We exemplary show the results from a public dataset of 981 patients and 8469 genes from the Cancer Genome Atlas (TCGA).

## I. Introduction

Machine learning and deep learning offer new opportunities to transform healthcare, and have been used in many different areas, including oncology [1], pathology [2], diabetes [3], or infectious diseases [4], [5]. However, clinical datasets are typically rather small and need to be aggregated over different hospitals. Often, data exchange over the internet is perceived as insurmountable, posing a roadblock hampering big data-based medical innovations. The most pressing problem in training powerful AI is that all the data usually distributed over various hospitals needs to be accessible in a central cloud. Due to recent data leaks, public opinion, and patient trust ([6], [7]) demand more secure ways of storing and processing data. As the E.U. GDPR (General Data Protection Regulations) and as the E.U. NISD (Network and Information Security Directive) entered into force in 2018 and 2016, respectively, data providers, researchers, and IT solution providers are challenged to find ways of providing hospitals complete control over how the patient data is processed. Federated AI enables collaborative AI without sharing the data and, thus, aligns with data protection regulations. Federated AI implies that each participant securely stores its data locally and only shares some intermediate parameters computed on local data [8], [9]. Graph Neural Networks (GNNs) are widely adopted within the biomedical domain [10]. Biological entities such as proteins do not function independently and thus must be analyzed on a systems level. GNNs provide a convenient way to model such interactions using e.g prior knowledge defined by a protein-protein interaction network (PPI). The PPI can be used as the GNN’s input graph, while the nodes can be enriched by patient-specific omics profiles. These knowledge-enriched deep learning models might be more interpretable compared to standard approaches when used to predict patient outcomes. However, due to the GDPR mentioned above, the challenge remains how to enable federated learning on Graph Neural Networks. Here, we present a software tool and Python package for federated ensemble-based learning with Graph Neural Networks. The implemented methodology enables federated learning by decomposing the input graph into relevant subgraphs based on which multiple GNN models are trained. The trained models are then shared by multiple parties to form a global, federated ensemble-based deep learning classifier.

## II. Materials and methods

### Input data

The input data for our software package consists of patient omics data on a gene level and a PPI network reflecting the interaction of the associated proteins. In order to perform graph classification using GNNs, each patient is represented by a PPI network, and its nodes are enriched by the patient’s individual omics features. We call these networks patient-specific PPI networks. Given that specific data representation, it is possible to classify patients based on their genomic characteristics while incorporating the knowledge about the functional relationships between proteins. It should be noted, that the topology of the network is the same for all patients.

### Ensemble learning with Graph Neural Networks

The proposed algorithm for graph-based ensemble learning consists of three steps:

1. Decomposition of the PPI network into relevance-weighted communities using explainable AI
2. Training of an ensemble GNN graph classifier based on the inferred communities
3. Predictions via Majority Voting

In the first step, the Python package GNNSubNet [11] is used to build a GNN classifier and to infer relevant PPI network communities (disease subnetworks). In detail, GNNSubNet utilizes the Graph Isomorphism Network (GIN) [12] to derive a graph classification model and implements a modification of the GNNExplainer [13] program such that it computes model-wide explanations. This is done by randomly sampling patient-specific networks while optimizing a single-node mask. From this node mask, edge relevance scores are computed and assigned as edge weights to the PPI network. A weighted community detection algorithm finally infers disease subnetworks. In the second step, an ensemble classifier based on the inferred disease subnetworks is created, and predictions are accommodated via Majority Voting. The ensemble members are predictive GNN models, that are based on the detected disease subnetworks, which overall makes the deep learning model more interpretable. High performing members of the ensembles may consist of a subnetwork biologically important for a specific disease or disease subtype.

In the *federated* case, each client has its dedicated data based on which GNN models of the ensemble are trained. These models are shared among all clients creating a global ensemble model, and predictions are again accomplished via Majority Vote (see Figure 1).

**Fig. 1.**
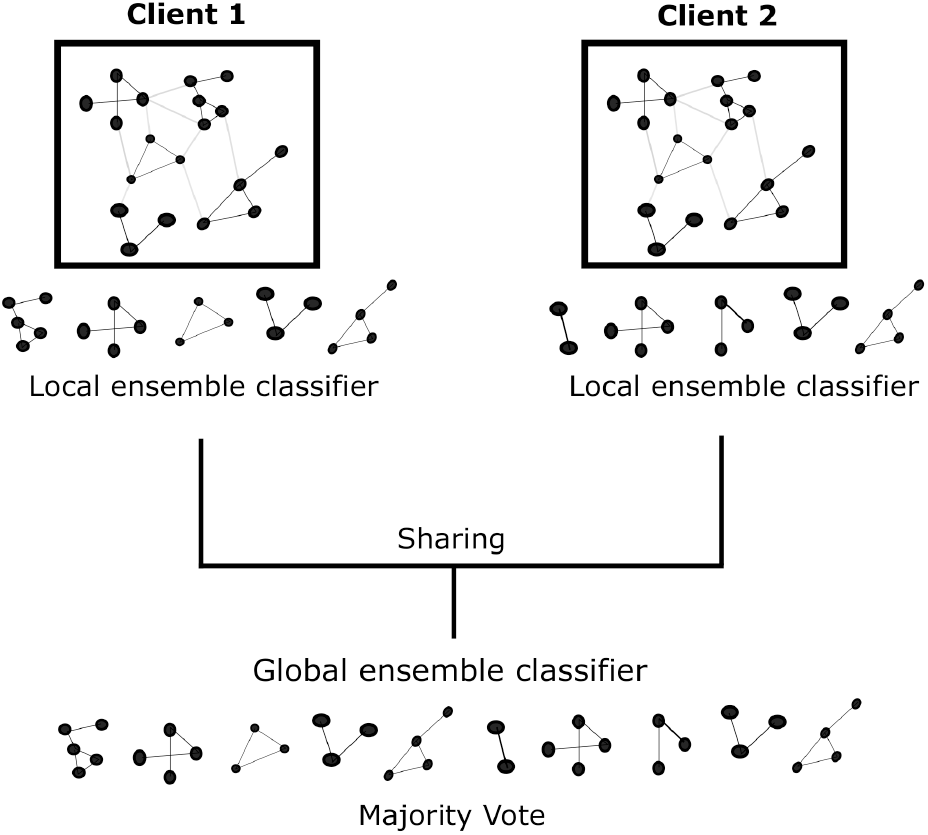
Federated Ensemble learning with Graph Neural Networks. Each client builds its dedicated ensemble classifier based on relevant subnetworks. The models trained on these subnetworks are shared and a global ensemble classifier is created. Final predictions are based on Majority Voting.

## III. Results and discussion

We used the gene expression data of human breast cancer patient samples for an experimental evaluation of the herein proposed methodologies. The Cancer Genome Atlas (TCGA) provided the data preprocessed as described in [14]. The data was structured by the topology of Human Protein Reference Database (HPRD) protein-protein interaction (PPI) network [15]. The resulting dataset comprises 981 patients and 8469 genes. The binary prediction task was to classify the samples into a group of patients with the luminal A subtype (499 samples) and patients with other breast cancer subtypes (482 samples).

### A. Performance of Ensemble-GNN in non-federated case

We assessed the performance of our algorithm using 10-fold cross-validation (see Table I). Ensemble-GNN, initially using GIN as a base learner, showed the average balanced accuracy of 0.86 (fourth row of Table I). As a comparison, a Random Forest (RF) classifier, not guided and restricted by any PPI knowledge graph, demonstrated 0.90 of average balanced accuracy on the same data set. The slight decrease in performance can be explained by the following reason: The GIN method [12] shows worse convergence during training on gene expression modality, compared to data with combined gene expression and DNA methylation modalities (see also [11], Table 2). Since transcriptomics is one of the most common omics types [16], we improved the performance of Ensemble-GNN specifically on gene expression modality using the ChebNet approach from [17]. ChebNet models have been successfully applied in [18] to classify patients based on gene expression profiles structured by a PPI network. The corresponding predictions were further explained on a patient level [18]. Ensemble-GNN employing ChebNet as a base learner achieved as good classification performance as RF (see Table I).

**TABLE I.** Performance within 10-fold CV (mean balanced accuracy ± standard error of the mean) of GNNSubNet and Ensemble-GNN utilizing TCGA-BRCA data, depending on a base learner (GIN or ChebNet).

| Method (base learner) | Balanced accuracy |
| --- | --- |
| RF (decision tree) | 0.90 ± 0.0109 |
| GNNSubNet (ChebNet) | 0.90 ± 0.0094 |
| Ensemble-GNN (ChebNet) | 0.90 ± 0.0105 |
| Ensemble-GNN (GIN) | 0.86 ± 0.0120 |
| GNNSubNet (GIN) | 0.52 ± 0.0121 |

**TABLE II.** Performance (mean balanced accuracy) of Ensemble-GNN based on TCGA-BRCA data, depending on the base learner GIN and ChebNet. For both federated setups, the data was distributed across three clients within five Monte-Carlo iterations.

| Method (base learner) | Federated setup with client-specific test data |  | Federated setup with global test data |  |
| --- | --- | --- | --- | --- |
|  | Local model | Federated model | Local model | Federated model |
| Ensemble-GNN (ChebNet) | 0.85 | 0.87 | 0.87 | 0.88 |
| Ensemble-GNN (GIN) | 0.81 | 0.85 | 0.78 | 0.83 |

### B. Performance of Ensemble-GNN in federated case

In the federated case, we evaluated the performance of Ensemble-GNN using the two implemented base learners: GIN and ChebNet. We utilized five Monte Carlo iterations, in which the data was equally distributed across three clients. Each client-specific data was then split into a train (80%) and test data set (20%). The trained models of the client-specific Ensemble-GNNs were combined into a global federated model. The performance of the global model was estimated using client-specific test sets.

For Ensemble-GNN (GIN), the mean local client-specific test accuracy from five Monte Carlo iterations was [0.84, 0.83, 0.78, 0.79, 0.81] with an overall mean value of 0.81 (see Table II). For the global, federated model we obtained [0.85, 0.86, 0.83, 0.84, 0.87] with a mean value of 0.85. Mean performance values for Ensemble-GNN using the ChebNet architecture were higher, 0.85 and 0.87 respectively. We have conducted an additional experiment, where we initially split the data into a global train set and a global test data set. The train set was then equally distributed across three clients. The average performance of the local classifier to predict the global test set was [0.78, 0.80, 0.77, 0.77, 0.76] with an overall mean value of 0.78. The accuracy of the federated model was [0.82, 0.83, 0.82, 0.80, 0.86] with an overall mean value of 0.83. The performance of Ensemble-GNN could be improved using ChebNet. In this case, we obtained mean values of 0.87 and 0.88 respectively (see Table II).

Note, we report on balanced accuracy in all cases to account for a possible unbalance sample distribution caused by the data splits.

## IV. Conclusion

We present Ensemble-GNN, a Python package for ensemble-based deep learning with interpretable disease sub-networks as ensemble members. The implemented methodology is especially suited, but not limited to, the federated case, where sensitive data is distributed across multiple locations. We could show that the models trained on subnetworks locally, and shared across multiple parties/clients globally, can improve client-specific predictive performance.

## V. Data availability

The Python package Ensemble-GNN including a comprehensive documentation of its usage is freely available from the GitHub repository https://github.com/pievos101/Ensemble-GNN.

